# Hierarchical Analysis of RNA Secondary Structures with Pseudoknots Based on Sections

**DOI:** 10.1101/2025.08.29.672997

**Authors:** Ryota Masuki, Donn Liew, Ee Hou Yong

**Affiliations:** Department of Applied Physics, The University of Tokyo, 7-3-1 Hongo, Bunkyo-ku, Tokyo 113-8656, Japan; Division of Physics and Applied Physics, School of Physical and Mathematical Sciences, Nanyang Technological University, Singapore 637371

## Abstract

Predicting RNA structures containing pseudoknots remains computationally challenging due to high processing costs and complexity. While standard methods for pseudoknot prediction require *O*(*N*^6^) time complexity, we present a hierarchical approach that significantly reduces computational cost while maintaining prediction accuracy. Our method analyzes RNA structures by dividing them into contiguous regions of unpaired bases derived from known secondary structures (“sections”). We examine pseudoknot interactions between sections using a nearest-neighbor energy model and dynamic programming. The algorithm scales as *O*(*n*^2^*ℓ*^4^), offering substantial computational advantages over existing global prediction methods. Our analysis of 726 transfer messenger RNA and 455 Ribonuclease P RNA sequences reveals that biologically relevant pseudoknots are highly concentrated among section pairs with large minimum free energy gains. Over 90% of connected section pairs appear within just the top 3% of section pairs ranked by MFE gain. For 2-clusters, our method achieves high prediction accuracy with sensitivity exceeding 0.90 and positive predictive value above 0.80. For 3-clusters, we discovered asymmetric behavior where “former” section pairs (formed early in the sequence) predict accurately, while “latter” section pairs show different formation patterns. The hierarchical section-based approach demonstrates that local energy considerations can effectively predict pseudoknot formations between unpaired regions. Our work provides strong evidence for the effectiveness of local energy calculations in pseudoknot prediction and offers insights into the dynamic processes governing RNA structure formation.

## Introduction

Ribonucleic acid (RNA) plays fundamental roles across all life forms, carrying genetic information, regulating gene expression, and performing various catalytic functions. Non-coding RNA, which does not translate into proteins, plays important roles in many catalytic and regulatory cellular processes^1–4^. The biological functions of non-coding RNA have strong connections with its molecular structure, particularly pseudoknots^5–8^. Thus, predicting and understanding the structure of RNA for any given base sequence is of great importance in biology, medicine, pharmacy, and other related fields. RNA secondary structure is defined as the set of base pairs that form a planar graph, excluding pseudoknots which are non-planar structural elements by definition^9–13^. RNA secondary structure forms the basic frame of an RNA structure, typically exhibiting greater thermodynamic stability than pseudoknot interactions. Several reliable algorithms have been developed to predict RNA secondary structure for any given base sequence, including dynamic programming algorithm (DPA)^14–19^ and stochastic methods^20–22^. Of the various numerical methods, the DPA method has proved to be one of the most popular and has been applied in many different areas^23, 24^. DPA accurately predicts RNA secondary structures by minimizing the free energy contribution under a local energy model in *O*(*L*^3^) time for an RNA sequence of length *L*.

Predicting RNA structures that include pseudoknots remains challenging due to computational complexity and high processing costs. The most straightforward approach would be to DPA to find minimum free energy structures across all possible base pairings. However, this approach is NP-complete even with simple energy models that only consider individual base pair contributions^25–29^. To address this computational challenge, researchers have developed algorithms that focus on specific subsets of pseudoknotted structures^30–35^. These methods improve efficiency by eliminating physically impossible configurations early in the analysis. For example, a dynamic programming algorithm achieving *O*(*N*^6^) time complexity and *O*(*N*^4^) space complexity was introduced^36^, providing the first method to fold optimal pseudoknotted RNAs using standard thermodynamic models. Though widely used, this approach becomes computationally intensive for longer RNA sequences. A later study showed that simpler dynamic programming techniques could achieve *O*(*N*^5^) time complexity^37^, offering improved performance but lacking the ability to handle complex pseudoknot configurations like kissing-hairpins, which are commonly found in RNA structures. Alternative approaches have emerged, including stochastic methods^38^ and novel computational strategies^39, 40^. While these techniques show promise in specific scenarios, comparative analyses^41^ reveal accuracy limitations, particularly when dealing with complex pseudoknot structures in RNA sequences longer than 200 nucleotides.

In order to overcome the trade-off between the accuracy of the model and the computational cost, there is now an increasing focus on hierarchical folding method^35, 42, 43^. The hierarchical folding method uses a two-step approach: first predicting the pseudoknot-free secondary structure, then calculating pseudoknot base pairs based on this foundation. A proposed DPA variant^44^ finds pseudoknotted structures in *O*(*N*^3^) time and *O*(*N*^2^) space by applying this hierarchical principle, matching the time efficiency of algorithms that only predict secondary structures. This approach can handle complex configurations like kissing hairpins, which traditionally required *O*(*N*^6^) time to process. However, comparative studies show these methods face significant challenges with accuracy, particularly for complex pseudoknots in longer sequences^31, 41, 45^. These limitations appear largely due to the fact that pseudoknot structures are often long-range and thus affected by the many possible 3D RNA conformations, which are difficult to account for in current models.

In this paper, we analyse local pseudoknot interactions by examining how different “sections” of an RNA (derived from known secondary structures) are connected via pseudoknots. Using a nearest-neighbor energy model and dynamic programming, we calculate the minimum free energy of potential pseudoknot formations between unpaired regions. Our approach focuses strictly on local energy contributions between sections and excludes interactions between different pseudoknots.

Building on this section-wise interaction framework, we develop a computationally efficient method that scales as *O*(*n*^2^*ℓ*^4^), where *n* represents the number of sections and *ℓ* represents their typical length. This hierarchical approach provides further understanding into how local structural elements contribute to global RNA configurations.

## Methods

### Definitions of terms

We define the terminologies used in the research of RNA structure.

1. **Primary structure** of an RNA refer to its chemical sequence.
2. **Secondary structure** of an RNA is the set of all local short-range pairing of complementary bases that can be represented in a planar graph without crossing lines. “Short-range” refers to non-crossing topological property and not the linear distance between bases in the sequence. Even pairings between distant bases (such as initial and final bases) are considered part of the secondary structure if they maintain this planar property. Common secondary structure motif includes hairpin loops, hairpin stems, helical duplexes, bulges, internal loops, multi-loops, and junctions.
3. **Pseudoknots** are long-range base pairings between secondary motifs that cannot be represented without crossing lines in a planar path. “Long-range” here refer to interactions that create crossing patterns when represented in a planar diagram and not the linear distance between paired bases in the sequence. These crossing interactions are tetiary structures. Pseudoknots are defined by these crossing patterns within the global RNA topology, not as isolated elements. The non-planarity emerges from the relationship between these interactions and the secondary structure, reflecting the topological complexity characteristic of pseudoknots.
4. **Pseudoknot base pair** is a base pair that participates in a non-nested pairing arrangement, creating crossing interactions when represented in a planar diagram. These pairs are part of the global topological structure where bases *i, j* form a pair and bases *k, l* form another pair such that *i* < *k* < *j* < *l*, resulting in crossing lines in a 2D representation.
5. **Pseudoknotted secondary structure** is defined as the set of all base pairs, including pseudoknots. It is specifically used when the set of base pairs includes pseudoknots.

The definitions above are explained graphically in Fig. 1a. Next, we will introduce some new terminologies that we will use for our analysis.

**Figure 1.**
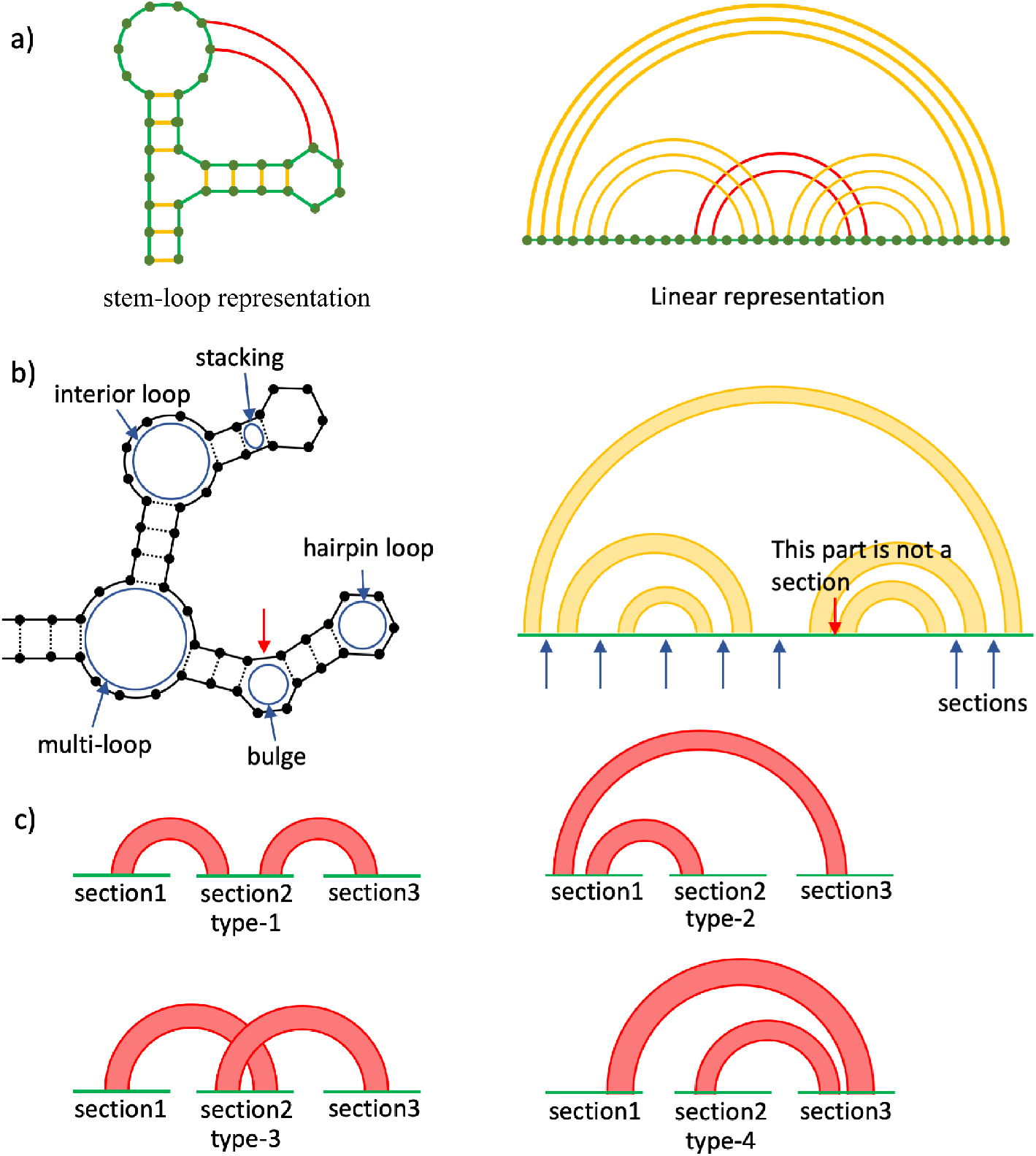
Graphical representations of RNA structures and section-based pseudoknot classifications used in this study. a) An example of RNA structure in stem-loop representation and linear representation. Yellow lines represent base pairs in the planar secondary structure that can be drawn without crossings. Red lines highlight the base pairs that create crossing interactions with the secondary structure, forming pseudoknot interactions. The pseudoknotted structure of this RNA is the set of all base pairs (both yellow and red lines). b) The stem-loop representation on the left and the linear representation on the right corresponds to the same structure. In the linear representation, the yellow band represents a set of consecutive base pairs. Each of the blue arrows in the linear representation points to a different section in the RNA sequence. The part that is pointed by the red arrow is not a section because the region does not include any unpaired bases. c) A list of all topologically distinct pseudoknot structures of 3-clusters.

1. A section is a contiguous subset of bases in the RNA sequence that are not involved in secondary structure base pairing, as determined from the known structures in our dataset. Specifically, if the *i*-th base and *j*-th base (*i* < *j*) are not paired in the secondary structure, they belong to the same section if all bases between them (from position *i* + 1 to *j* −1) are also not involved in secondary structure base pairing. Each section represents a single-stranded region that may potentially form pseudoknot interactions with other sections. This definition relies on having accurate secondary structure information, which in our study comes from established RNA structure databases.
2. **Section pair** refers to a pair of sections. The *i*-th section and *j*-th section are **connected** if there exists a pseudoknot that connects a base in *i*-th section and another in *j*-th section. See Fig. 1b for a graphical description of sections.
3. *N***-cluster** is a cluster consisting of *N* sections connected by pseudoknots. The number of topologically distinct pseudoknot structures increases rapidly once *N ≥* 3. We will only consider 2-clusters and 3-clusters in this work. The different types of 3-clusters are shown in Fig. 1c.

### Energy model

We use the nearest-neighbor energy model which is based on the chemical and thermodynamic properties of the RNA chains. Nearest-neighbor energy model is widely used to predict secondary structure in conjunction with DPA^46^–53. In the nearest-neighbor energy model, values of free energy contribution are assigned to each closed loop (e.g. stackings, bulge loops, interior loops, multi-loops) in RNA secondary structure, and the total free energy of the structure is calculated as the sum of all the free energy contribution from the loops^54^. The parameters and detailed rules in the energy model used in our analysis are based on mfold version 3.6^15, 16^. While these parameters were derived from experiments in salt-free conditions, we apply them as an approximation for pseudoknot prediction in the absence of comprehensive energy tables calibrated for varying salt concentrations.

Formally, the energy model is defined as

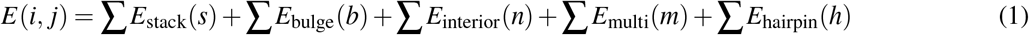

Using equation 1, we calculate the free energy contribution of the pseudoknot between two sections in the example shown (in linear representation) in Fig. 2a. According to the conventional rule of the nearest-neighbor energy model, the free energy contribution of the pseudoknot in Figure 2a is given by:

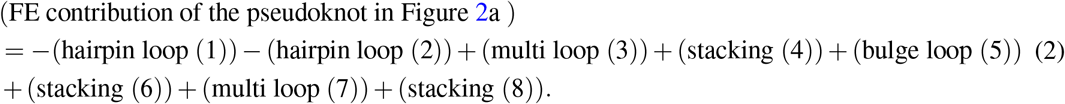

**Figure 2.**
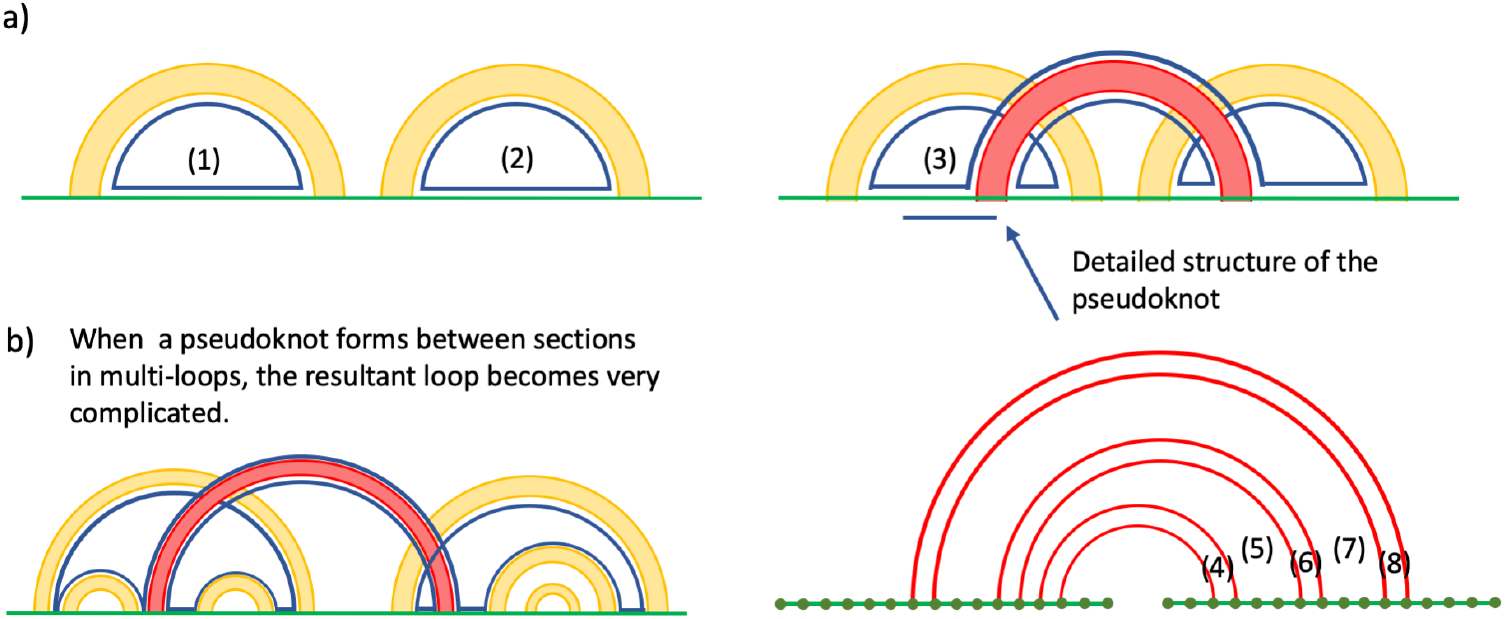
Energy model calculation examples demonstrating local versus global structural considerations. a) An example of base pair structure of a pseudoknot. The yellow bands represent sets of base pairs belonging to a secondary structure, while the red band represents a set of consecutive pseudoknots. The figure in the bottom right shows the detailed base pair structure in the pseudoknot. b) An example of base pair structure of a pseudoknot between sections in multi-loops. The newly formed loop that includes the pseudoknot contains all the sections in the multi-loops that the pseudoknot connects, illustrating the computational complexity of global energy calculations.

However, in this work, the contribution of loops that include base pairs in secondary structures are neglected, and only loops that include pseudoknot base pairs are considered. Hence, the free energy contribution of the psuedoknot in Figure 2 is calculated as

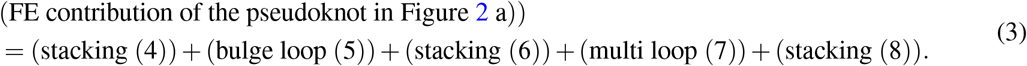

Our energy model uses two main nearest-neighbour thermodynamic parameters: stacking energies and loop energies. Stacking energies represent free energy contributions of stacked pairs formed by consecutive base pairs, with values ranging from -3.4 kcal/mol (most stable) to +1.5 kcal/mol (least stable). Loop energies compose of multiple parameter arrays such as destabilising values for interior loops (4.1 to kcal/mol for lengths 1–30), bulge loops energies (3.9 to 6.7 kcal/mol), mismatch energy parameters (-2.7 to +0.2 kcal/mol)^16^. Unlike standard models that calculate global free energy, our approach focuses exclusively on local pseudoknot interactions between RNA sections. By ignoring energy contributions from base pair loops and applying calculations independently to each section pair, we enable efficient hierarchical structure prediction. For exact values of our thermodynamic parameters, please refer to our code.

We exclude the free energy contribution from loops containing secondary structure base pairs to focus on local interactions between sections under analysis. Including such loops would contradict our hierarchical approach to treat pseudoknot formations independently. This exclusion is necessary, since including these energy terms would require information about the global RNA configuration.

Specifically, when sections within our target pair reside in a multi-loop (Fig. 2b), calculating their local pseudoknot structure would necessitate predicting pseudoknot formations across other section pairs simultaneously. This would force us to expand from local to global structure prediction, defeating the purpose of our section-based approach. However, the global base pair structure predicted by DPA method is known to be inaccurate even when additional energy parameters for certain topological motifs of pseudoknots are included^44^. This is most probably due to the inability to account for 3D conformation effectively even though the pseudoknot structure is strongly affected by the 3D RNA structure^55, 56^. In this work, instead of trying to incorporate the effects of 3D conformation into the DPA method, we focused on predicting local structure using only local information.

Our energy model simplifies the entropic contribution to total free energy by treating loop contributions as purely additive. This approximation for pseudoknotted structures enables computational efficiency while maintaining our focus on local interactions. By analyzing section pairs independently and excluding loops containing pseudoknot-free base pairs, we achieve a computational cost of *O*(*n*^2^*ℓ*^4^). The empirical validity of this approximation is demonstrated in our Results section.

While our approach for energy calculations shares mathematical similarities with RNA hybridization algorithms through nearest-neighbor dynamic programming models, there are key differences in our implementation. Our method specifically addresses intramolecular interactions between sections within a single RNA molecule, unlike hybridization algorithms that focus on intermolecular binding^57, 58^. Our approach also incorporates topological constraints specific to pseudoknot formation within an existing RNA secondary structure. For 3-clusters analysis, we apply weighted energy minimization (as shown in Fig. 6, where *w* = 0.8 proved optimal), a consideration not typically present in standard hybridization algorithms^59^–61. Furthermore, our method distinguishes between different pseudoknot topologies (Fig. 1c) and prevents crossing pseudoknot interactions within section pairs. We apply these techniques to analyse local pseudoknot interactions within known unpaired regions, which helps demonstrate how RNA structures can be examined in a hierarchical manner.

### Algorithm

In this subsection, we present the algorithm to determine the MFE structure between section pair. The MFE structure between section pair is calculated by considering all possible structures where base pairs can form pseudoknots with the secondary structure, but with the constraint that pseudoknot base pairs within the same section pair cannot cross each other. This allows pseudoknots between different sections while preventing nested pseudoknots within the same section pair. Moreover, two sections from the same loop is prohibited because the base pairs between these sections will be in secondary structures, rather than pseudoknots. Sections are identified by analyzing the loop regions from the predicted secondary structure, where each continuous stretch of unpaired nucleotides within a loop forms a distinct section. The algorithm systematically identifies all sections by scanning the RNA sequence and grouping consecutive unpaired bases that belong to the same loop structure.

We denote the two sections in the given section pair as “section1” and “section2”, with length *ℓ*_1_ and *ℓ*_2_ respectively. We define a matrix *C* whose entry *C*(*i, j*) (1 ≤ *i* ≤ *ℓ*_1_, 1 ≤ *j* ≤ *ℓ*_2_) is defined as the minimum value of free energy contribution over candidate structures in which *i*th base in section1 and *j*th base in section2 are paired and *i*^*′*^th base in section1 or *j*^*′*^th base in section2 are not connected when either *i*^*′*^ < *i* or *j* < *j*^*′*^ We set *C*(*i, j*) = ∞ when the *i*th base in section1 and the *j*th base in section2 cannot form a base pair. By definition, the MFE between the section-pairs that we are calculating is the minimum entry of the *C* matrix. Thus, the problem of finding the MFE turns into the calculation of matrix *C*. The entries of matrix C can be calculated using the following recursion relation.

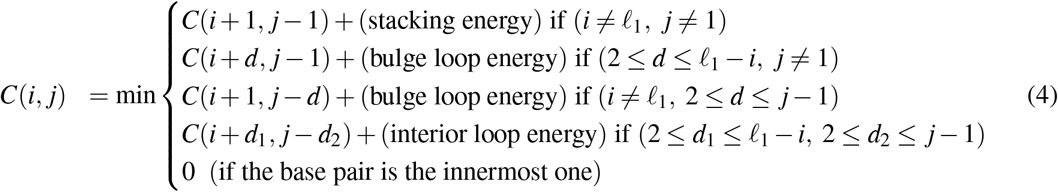

After calculating the *C* matrix (see Algorithm 1), we determine the MFE structure between section pairs by first selecting the binding sites. Binding sites are selected through dynamic programming that minimizes free energy, where the algorithm identifies the minimum entry in the *C* matrix, representing the optimal base pairing configuration. Base pairs are formed only between complementary bases (AU, UA, CG, GC, GU, UG). For sections to interact, they must be from different loops, contain complementary bases and have negative free energy contribution due to their interaction. For 2-clusters, we use the MFE structure between the section pair. For 3-clusters, we apply weighted energy minimization: (energy of former pair) + *w*× (energy of latter pair), where 0 < *w* ≤ 1. We ensure consistency by preventing bases already paired in former sections from forming pairs in latter sections. This hierarchical approach constructs the global structure by identifying favorable local interactions first, then combining them while maintaining structural constraints.

We provide a proof for the computational complexity of our algorithm. If *n* is the number of sections and *l* the typical length of one section, we compute the MFE for all possible section pairs for *n* sections, yielding *n*(*n* − 1)*/*2≈ *O*(*n*^2^). For each section pair with lengths *l*_1_ and *l*_2_, we construct matrix *C* of size *l*_1_ × *l*_2_. For each entry *C*(*i, j*), there are three cases: stacking with *O*(1) complexity, bulge loop with *O*(*l*_1_) and *O*(*l*_2_) complexity respectively, interior loop with *O*(*l*_1_ × *l*_2_) complexity due to two nested loops of length *l*_1_ and *l*_2_. Thus, the worst-case time complexity for each *C*(*i, j*) entry is *O*(*l*_1_ × *l*_2_ × *l*_1_ × *l*_2_) or *O*(*l*^4^) assuming *l*_1_ ≈ *l*_2_ ≈ *l*. Thus, for all *O*(*n*^2^) section pairs, the total computational cost is *O*(*n*^2^*l*^4^). For typical RNA molecules in our dataset, we observed *n* ≈ *N/l*, where *N* is the total RNA length. This relationship suggests our algorithm’s complexity in terms of total RNA length is approximately *O*(*N*^2^).

## Results and Discussion

In this work, 726 transfer messenger RNA sequences and 455 Ribonuclease P RNA sequences from the RNAstrand database^62^ were analyzed. These RNA families were selected because they represent a comprehensive and diverse dataset of biologically relevant structures with well-documented pseudoknots. Rather than selecting a small subset of examples, we analyzed the complete set of these RNA families available in the database to ensure a thorough and unbiased evaluation of our approach. Both families consist of relatively long RNA sequences with many pseudoknots, making them ideal test cases for our section-based prediction method.

### Basic Properties of investigated RNA sequences

Table 1 summarises key characteristics of transfer messenger RNA and ribonuclease P RNA sequences from the RNAstrand database. Both RNA classes show remarkable structural similarities, with average sequence lengths of approximately 350 nucleotides, around 30 sections per sequence, and average section lengths of about 6 nucleotides. Transfer messenger RNA sequences show greater variability in their measurements. The increase in standard deviation is due to sequences exceeding 1000 bases, with each sequence containing a single extended section that forms no pseudoknot connections with other sections. These outlier sequences do not impact our analysis of pseudoknot interactions between section pairs. The substantial length of these RNA sequences presents a particular challenge, as conventional prediction methods struggle to accurately determine the structure of such extended sequences^31^.

**Table 1.** Structural characteristics of RNA sequences analyzed from RNAstrand database.

| Parameter | Transfer messenger RNA (TMR) | Ribonuclease P RNA (RPA) |
| --- | --- | --- |
| Sequences analyzed (n) | 726 | 455 |
| Mean sequence length (nt) | 367.7 | 332.5 |
| SD sequence length | 86.3 | 49.5 |
| Mean sections per sequence | 29.8 | 28.1 |
| Mean section length (nt) | 7.7 | 5.2 |
| SD section length | 23.1 | 6.1 |

### Predicting the connected section pairs based on MFE

In this section, we present our analyses based on section pairs. The histogram of MFE gain (absolute value of MFE) for all section pairs in the investigated database is shown in Figure 4a. As the MFE gain increases, the corresponding frequency shows an exponential decay.

**Figure 3.**
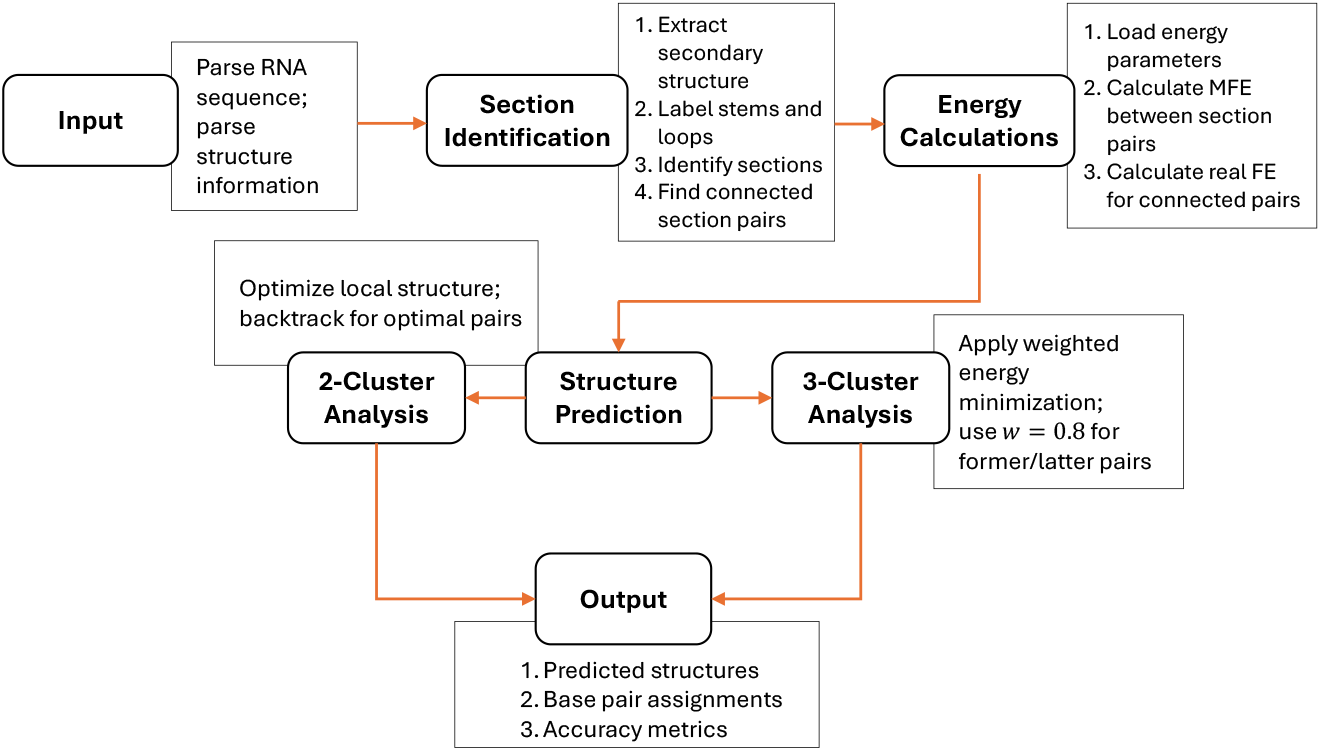
Implementation flowchart of the RNA pseudoknot structure prediction algorithm. The flowchart illustrates the hierarchical approach of our method, starting with input parsing, followed by section identification from the secondary structure. Energy calculations determine the minimum free energy (MFE) between section pairs. Structure prediction then branches into 2-cluster and 3-cluster analysis paths, with the latter using weighted energy minimization (*w* = 0.8) to balance former and latter section pair contributions. The output includes predicted structures, base pair assignments, and accuracy metrics.

**Figure 4.**
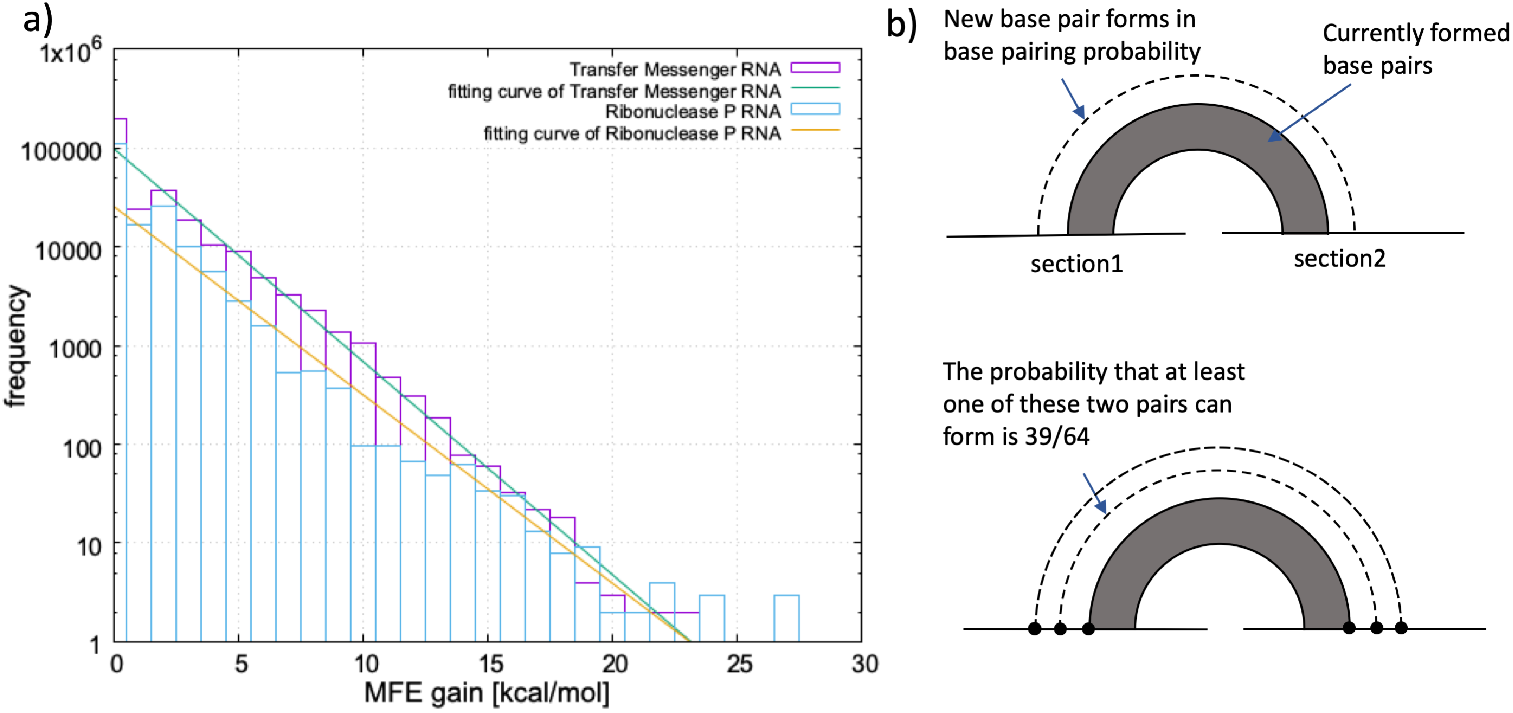
Distribution of MFE gain for section pairs in RNA databases. a) Histogram showing exponential decay of MFE gain frequency for all section pairs in transfer messenger RNA and Ribonuclease P RNA sequences from RNAstrand database. b) Simple probabilistic model explaining the observed exponential decay, assuming independent base pairing probability and 1.0 kcal/mol energy contribution per base pair.

The exponential decay pattern (Fig.4a) can be explained using a simple model (Fig.4b). Our model makes two key assumptions: (1) When pseudoknot base pairs already exist between two sections, the probability of forming another base pair between these same sections is independent of the existing structure. We call this the “probability of base pairing”. (2) Each base pair contributes approximately -1.0 kcal/mol to the free energy, which represents the average stacking energy across all possible stacking patterns in our energy model. With these two assumptions, the histogram in Fig. 4a can be fitted to the exponential function with fitting parameter *a*:

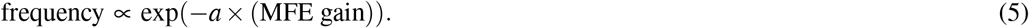

#### Algorithm 1 Dynamic Programming Algorithm for MFE calculation between section pairs

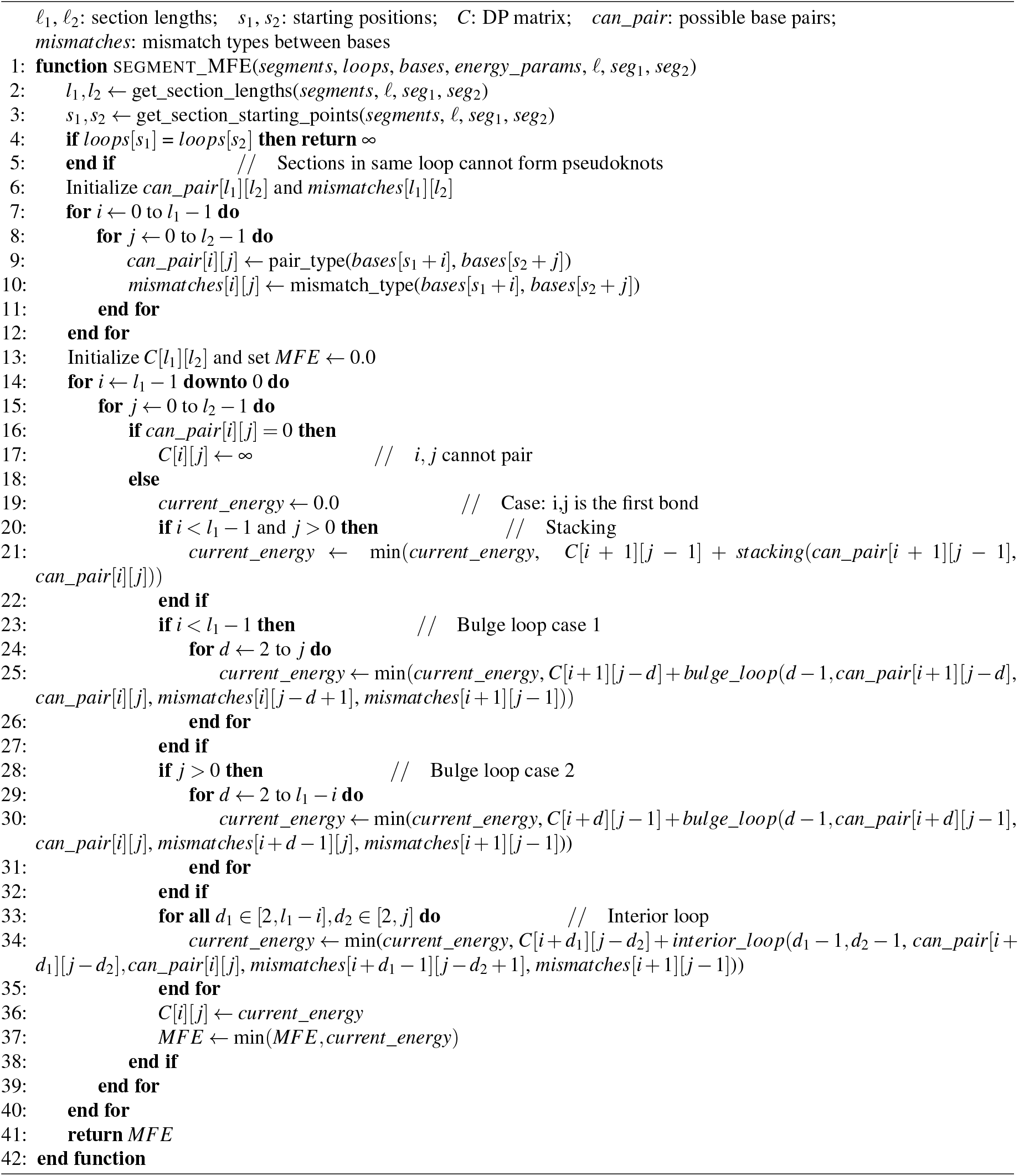

The probability of base pairing can be calculated as

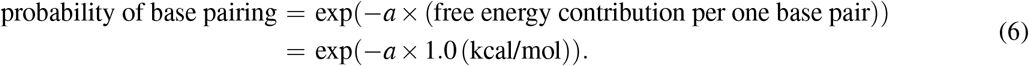

Curve fitting analysis reveals base pairing probabilities of 0.608 for transfer messenger RNA and 0.632 for ribonuclease P RNA. These similar probability values can be understood by considering the basic constraints of RNA base pairing. In sections with random base sequences, the probability of two adjacent bases forming a new base pair is 3/8 (0.375), as there are six possible pairing types (AU, UA, CG, GC, UG, GU). The probability of at least one of two adjacent base pairs forming next to an existing pair is 1 − (5*/*8)^2^ = 39/64 (0.609), which closely matches our observed pairing probabilities.

Our results that MFE structures between section pairs primarily consist of stacking interactions and small internal loops of length 2. The exponential decay in the distribution means that section pairs with large MFE gains are relatively uncommon. Specifically, only 5.5% of pseudoknots in transfer messenger RNA and 2.8% in ribonuclease P RNA have MFE gains exceeding 5.0 kcal/mol. Even fewer exceed 10.0 kcal/mol: just 0.47% in transfer messenger RNA and 0.23% in ribonuclease P RNA.

We investigated the connecting probability for both transfer messenger RNA and ribonuclease P RNA in our database (Fig. 5). Connecting probability is defined as the ratio between connected section pairs and all section pairs with a given minimum free energy:

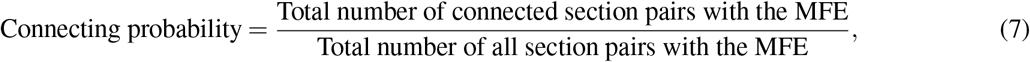

**Figure 5.**
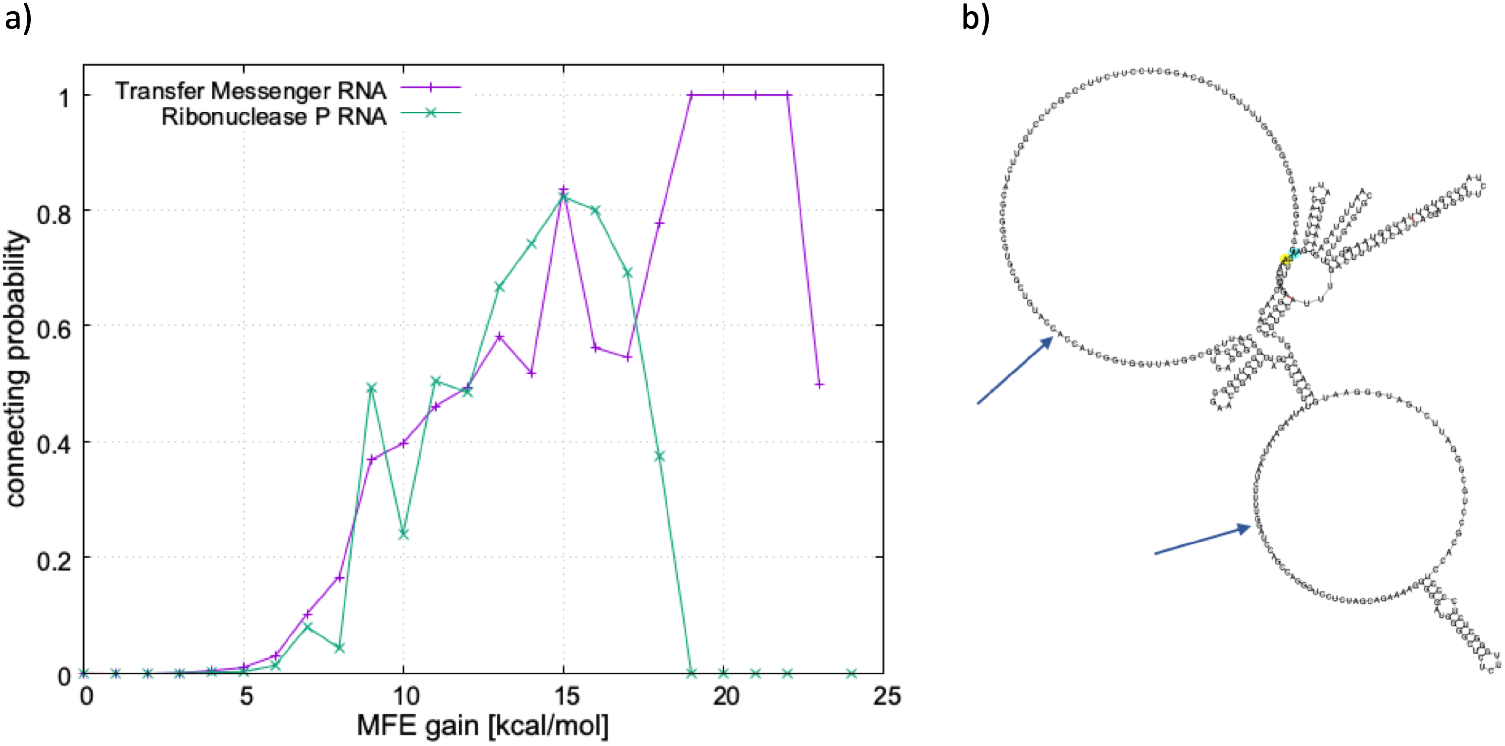
Analysis of pseudoknot formation likelihood based on energetic predictions. a) Connecting probability as a function of MFE gain for transfer messenger RNA and ribonuclease P RNA sequences, defined as the ratio of actually connected section pairs to all section pairs with a given MFE value. b) Representative example of a non-connecting section pair with substantial MFE gain (80 kcal/mol), illustrating how conformational entropy losses can outweigh local energy gains in long sections (104 and 48 nucleotides).

**Figure 6.**
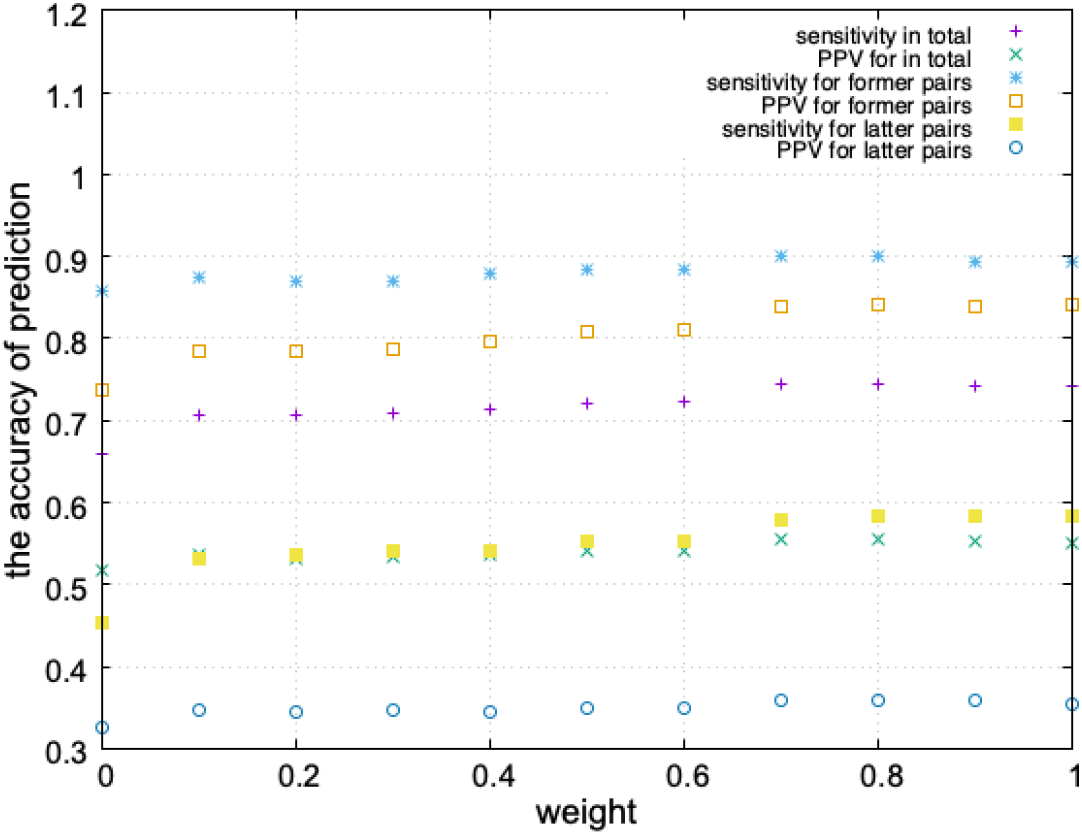
Optimization of weighting parameter for 3-cluster pseudoknot structure prediction. Prediction accuracy metrics (sensitivity and positive predictive value) as a function of weight parameter w applied to the free energy contribution of latter section pairs in the weighted energy minimization approach. Optimal performance occurs at w = 0.8, indicating preferential weighting of former section pair interactions while maintaining contribution from latter pairs in type-1 3-cluster configurations.

We expected a positive correlation between MFE gain and connecting probability, since larger free energy gains typically create more stable structures. However, we observed non-connecting section pairs with unexpectedly large MFE gains. Despite the large absolute MFE gain, these sections have relatively small free energy gain per base pair compared to genuinely connected section pairs. Long non-connecting sections with a large total MFE gain but a much smaller gain per base pair is due to the inclusion of numerous paired bases in MFE structures. The loss of conformational entropy, which our nearest-neighbor energy model ignores, outweighs the stabilising effect of each individual pair.

From the analyses above, we see that a large proportion of connected section pairs are those with large MFE gain, which account for a small percentage of total section pairs. All the section pairs in the investigated database were sorted in ascending order in terms of MFE gain, and the relation between cumulative frequencies in all section pairs and those in connected section pairs was investigated for all connected section pairs. We find that more than 90% of connected section pairs in transfer messenger RNA are included in the top 10^4^ section pairs (3%) with largest maximum free energy gains (out of all the 3.5 × 10^5^ possible section pairs). For the case of ribonuclease P RNA, more than 90% of connected section pairs are included in the top 1500 section pairs (1%) with largest maximum free energy gains (out of all the 1.8 × 10^5^ possible section pairs). Therefore, it is sufficient to only consider a small fraction of section pairs with large MFE gain in order to predict RNA structures with pseudoknots. Our analysis reveals that connected section pairs predominantly appear among those with large MFE gains. Upon sorting all section pairs by MFE gain, we found a concentration of biologically relevant connections. In transfer messenger RNA, over 90% of all connected section pairs appear within just the top 10^4^ section pairs (3% of the 3.5 × 10^5^ total possible pairs) ranked by maximum free energy gain. The concentration is even more pronounced in ribonuclease P RNA, where 90% of connected pairs are found within the top 1,500 section pairs (just 1% of 1.8× 10^5^ possible pairs). On average, we need to consider only about 14 section pairs per transfer messenger RNA sequence (10,000/726) or 3 section pairs per ribonuclease P RNA sequence (1,500/455) to account for 90% of pseudoknots. This reduces the computational search space when predicting RNA structures with pseudoknots, enabling more efficient algorithms by focusing exclusively on section pairs with the largest MFE gains.

### Comparison between MFE Structures and Real Structures for Connected Section Pairs

Our dataset analysis reveals that connected section pairs exclusively form either 2-clusters or 3-clusters, with no observed instances of larger cluster formations (Table 2). 2-clusters predominate in both RNA classes, while 3-clusters occur rarely. Due to their distinct properties, we analyse these cluster types separately. For the 2-clusters in our database, we compared the MFE of connected section pairs with the free energy contribution of their actual structure (real FE). While the proportion of 2-clusters where real FE equals MFE varies considerably between RNA classes (68.7% for transfer messenger RNA versus 39.9% for ribonuclease P RNA), we note that more than 80% of all 2-clusters demonstrate a free energy gain exceeding 0.8× MFE gain. This pattern suggests energetic principles governing 2-cluster formation across different RNA classes.

**Table 2.** The comparison of MFE and free energy contribution of real structure for connected section pairs in 2-clusters.

| Parameter | TMR | RPA |
| --- | --- | --- |
| The number of all 2-clusters | 2337 | 534 |
| The number of all 3-clusters | 110 | 1 |
| The number of 2-clusters for which MFE = real FE | 1605(68.7%) | 213(39.9%) |
| The number of 2-clusters for which real FE < $0.8 \times$ MFE | 1972(84.4%) | 438(82.0%) |
| Sensitivity of base pair prediction for 2-clusters | 0.9159 | 0.9087 |
| PPV of base pair prediction for 2-clusters | 0.8400 | 0.8056 |

Given the strong correlation between minimum free energy (MFE) configurations and actual structures in 2-clusters, we conducted a comparison of their base pair arrangements. For each 2-cluster across both RNA databases, we uses three standard metrics: true positives (TP, correctly predicted base pairs), false positives (FP, predicted base pairs absent in actual structures), and false negatives (FN, actual base pairs missed by our predictions) to quantify prediction accuracy. When multiple MFE structures existed for a single 2-cluster, we calculated the average values for these metrics to ensure equal weighting across all clusters. From these measurements, we calculate sensitivity and positive predictive value comprehensive (PPV):

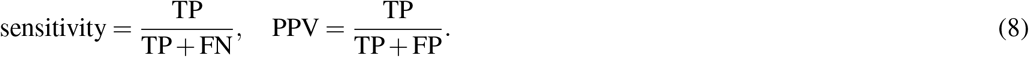

The sensitivity is over 0.9 and PPV is over 0.8 for both RNA classes, which shows that the base pair structure of 2-clusters can be well predicted using the simple MFE calculation. These accuracy values significantly outperform previous global prediction methods, which typically achieve sensitivity of 0.5-0.7 and PPV below 0.7 for complex pseudoknotted structures^41^. This improvement demonstrates the advantage of our local section-based approach over global structure prediction methods. The high accuracy of our predictions suggests pseudoknot structures are predominantly determined by local properties rather than global context. The locality property supports a hierarchical approach to RNA structure prediction. In other words, local pseudoknot prediction can be effectively decoupled from global structure determination. With our earlier observation that only a small fraction of high-MFE section pairs need consideration, the hierarchical approach offers potential for analysing longer RNA sequences. Thus, more sophisticated computational models can be applied to complex RNA structures.

It is important to acknowledge that our analysis relies on knowledge of the true secondary structure from which we derive the unpaired sections. This differs from the more challenging problem of *de novo* pseudoknot prediction without prior structural information. While our results demonstrate that local energetic considerations can effectively predict pseudoknot formations between known unpaired regions, additional research would be needed to determine whether similar accuracy could be achieved when starting with predicted rather than true secondary structures.

To further examine pseudoknot patterns in our dataset, we classified them according to their structure. Unlike 2-clusters, 3-clusters exhibit several topologically distinct configurations (Fig. 1c). Our dataset contains 110 type-1 3-clusters in transfer messenger RNA and only a single type-2 3-cluster in ribonuclease P RNA. This distribution demonstrates that 2-clusters are significantly more prevalent than 3-clusters in RNA structures. Given the limited sample size of type-2 3-clusters, our analysis focuses exclusively on the 110 type-1 3-clusters found in transfer messenger RNA. Within these type-1 3-clusters, we designate the connection between the first two sections (section1 and section2 in Fig. 1c) as the “former section pair”, and the connection between the second and third sections (section2 and section3) as the “latter section pair”.

We analysed the 110 type-1 3-clusters in transfer messenger RNA by comparing MFE calculations with actual structures for both former and latter section pairs. Our methodology treated these pairs independently, using the same approach applied to 2-clusters without considering that section2 participates in both connections within the 3-cluster structure. For former section pairs, approximately 50% have real structures identical to predicted MFE structures, while over 80% have free energy contributions lower than 0.8× MFE (Table 3). These results closely resemble our 2-cluster prediction accuracy. In contrast, latter section pairs show markedly different behavior, with only a small fraction having free energy contributions approximating their MFE. The sensitivity and PPV metrics further highlight this disparity. Former section pairs demonstrate prediction accuracy comparable to 2-clusters, while latter section pairs exhibit particularly low PPV values. Across both pair types, sensitivity consistently exceeds PPV, indicating our method tends to predict excess base pairs. This over-prediction is expected, as our approach treats these connections as independent 2-clusters despite their presence in more complex 3-cluster structures.

**Table 3.** The comparison of MFE and free energy contribution of real structure for connected section pairs in 3-clusters.

| Parameter | Former section pair | Latter section pair |
| --- | --- | --- |
| Total number | 110 | 110 |
| The number of section pair for which real FE = MFE | 55(50.0%) | 10(9.1%) |
| The number of section pair for which real FE < 0.8 × MFE | 91(82.7%) | 18(16.3%) |
| Sensitivity of base pair prediction | 0.8343 | 0.5529 |
| PPV of base pair prediction | 0.7164 | 0.2568 |

While former section pairs (between sections 1-2) can be predicted with accuracy comparable to 2-clusters using simple MFE calculations, this approach fails for latter section pairs (between sections 2-3). The failure suggests that former section pairs form independently, whereas latter section pair formations are strongly influenced by pre-existing base-pair connections. This asymmetry appears to reflect the fundamental biology of RNA synthesis, which proceeds directionally from 5’ to 3’ along the sequence. During transcription, former section pairs likely establish their interactions before latter sections are even fully synthesised, as supported by previous research on co-transcriptional folding dynamics^63, 64^. The consistency of this pattern across our dataset suggests it represents a general feature of RNA folding rather than a family-specific characteristic. These findings challenge the assumption that RNA structures in cells form solely through global free energy minimization^65^–68. Instead, our results suggest that the dynamic and sequential nature of RNA synthesis/folding plays a crucial role in determining the final structural configuration, particularly for complex pseudoknot arrangements.

### The Comparison between MFE Structures and Real Structures for 3-clusters

Our previous analysis treated section pairs independently, overlooking the constraint that a single base in section2 cannot simultaneously pair with bases in both section1 and section3. This section presents our approach to predicting complete 3-cluster structures using the nearest-neighbor energy model while considering this constraint. Based on our findings regarding the asymmetry between former and latter section pairs, we developed two prediction strategies:

1. Equal weighting: Minimize the combined free energy contribution from both former and latter section pairs, treating them with equal importance.
2. Sequential determination: First identify the minimum free energy structure of the former section pair, then determine the latter section pair structure while ensuring no bases in section2 are reused.

These approaches represent special cases of a more general weighted method for predicting 3-cluster pseudoknot structures. This method minimizes:

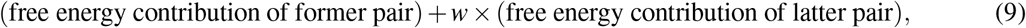

where *w* is the weight of free energy contribution of latter section pair compared to that of former section pair, that takes a value in the range 0 < *w* ≤ 1. We implemented an energy minimisation algorithm that evaluates all possible base pairings between sections (Figure 6). For each combination of former section pairs (sections 1-2) and latter section pairs (sections 2-3), we calculate their respective minimum free energy contributions using nearest-neighbor thermodynamic parameters based on the energy model prescribed by mfold version 3.6. The algorithm explores all possible base pairing configurations through dynamic programming, considering stacking interactions, bulge loops, and interior loops. To determine the optimal weight parameter (*w*), we evaluate multiple values between 0 and 1, calculating for each: (1) the total weighted energy for all possible section pair combinations, (2) the corresponding base pair predictions, and (3) the accuracy metrics (sensitivity and PPV) by comparing against the known structures. The computational implementation ensures that bases already paired in the former section pair cannot participate in the latter section pair’s interactions, maintaining the physical constraints of RNA folding. This comprehensive sampling across the weight parameter space revealed that *w* = 0.8 provides optimal prediction accuracy, suggesting a natural bias toward former section pair interactions while still accounting for latter pair contributions. However, the prediction of pseudoknot structure in latter section pairs is far less accurate than those in former section pairs for all value of *w*. This suggests the pseudoknot structure of latter section pair in 3-clusters is influenced by the global structure or 3D conformation of the RNA, that are beyond the limit of this energy model based on local substructures.

A limitation of our approach is the use of energy parameters derived from salt-free experimental conditions (mfold 3.6) to predict structures that typically form in the presence of salt. This approximation was necessary due to the lack of comprehensive energy tables for RNA in various salt concentrations. Despite this limitation, our method demonstrates strong predictive power for 2-clusters, suggesting that local pseudoknot formation may be reasonably approximated by these parameters even without explicitly modeling ionic effects. Future work could significantly improve prediction accuracy by incorporating salt-dependent energy parameters as they become available.

Our method is successful despite using additive entropy approximation for pseudoknots. Although previous work has shown that entropy contributions in pseudoknotted structures are not strictly additive^69, 70^, our results for 2-clusters (sensitivity *>* 0.90, PPV *>* 0.80) suggest this simplification works well within our hierarchical section-based framework. The high accuracy indicates that for common pseudoknot configurations, local energy minimization with simplified entropy terms captures the essential physics governing structure formation. The method’s limitations with latter section pairs in 3-clusters likely indicate where more complex entropic considerations become necessary. Nonetheless, this simplification allows for structural prediction of longer RNA sequences while maintaining good accuracy for the most common pseudoknot configurations.

## Conclusion

We have presented a section-based approach to RNA pseudoknot analysis that employs mfold nearest-neighbor energy model with DPA. This method offers both computational efficiency and prediction accuracy advantages over traditional approaches. Our algorithm scales as *O*(*n*^2^*ℓ*^4^), where *n* represents the number of sections and *ℓ* represents the average section length. Assuming section length remains independent of total RNA sequence length, the computational complexity reduces to *O*(*L*^2^) for an RNA of length *L*. A key finding of our research is the highly concentrated distribution of biologically relevant pseudoknots among section pairs with large MFE gains. By focusing on just the top 3% of section pairs ranked by MFE gain, we can capture approximately 90% of all connected section pairs. This observation enables substantial reductions in the search space required for structure prediction.

Our approach predicts 2-cluster pseudoknot structures with good accuracy, achieving sensitivity above 0.90 and positive predictive value exceeding 0.80. These results significantly outperform conventional methods that attempt to predict global pseudoknotted structures in a single step, demonstrating the effectiveness of our hierarchical, section-based strategy. This indicates that once unpaired regions are identified, 2-cluster pseudoknot formations between them appear to be primarily determined by local information. This observation suggests potential value in hierarchical approaches to RNA structure prediction, though such approaches would need to address the challenge of accurately identifying relevant unpaired regions and resolving potential competition between pseudoknot and regular secondary structure formation. Our work provides evidence for the effectiveness of local energy calculations specifically for the subproblem of pseudoknot prediction between known unpaired regions. Despite our method’s effectiveness for 2-cluster, there is significant limitations when applied to 3-clusters. Nearly all 3-clusters conform to the type-1 configuration (Fig. 1c). While our approach can predict former section pairs in these 3-clusters with accuracy comparable to 2-clusters, it performs poorly when predicting latter section pairs. The inadequacy of our local energy model for latter section pairs suggests that these pseudoknots are significantly influenced by factors beyond local interactions, such as the dynamic process of RNA folding and the global 3D RNA conformation. These findings highlight the limitations of purely thermodynamic approaches to complex RNA structure prediction and suggest that accurate modeling of 3-clusters may require incorporating kinetic folding pathways and long-range structural constraints.

## Acknowledgements

R.M. acknowledges support by the GRI programme from Nanyang Technological University and the SVAP program from The University of Tokyo. D.L. and E.H.Y. acknowledges support from Nanyang Technological University, Singapore, under its Start Up Grant Scheme (04INS000175C230), Singapore Ministry of Education through the Academic Research Fund Tier 1 (RG140/22) and Academic Research Fund Tier 2 (MOE-T2EP50223-0014). The authors thank Dr. Michael Zuker for his permission to use the mfold energy model that he developed.

## Author contributions statement

R.M. and E.H.Y. conceived the project; R.M. carried out the simulations; R.M., D.L. and E.H.Y. wrote the paper.

